# Temporally and Spatially Partitioned Neuropeptide Release from Individual Clock Neurons

**DOI:** 10.1101/2020.07.24.219725

**Authors:** Markus K. Klose, Marcel P. Bruchez, David L. Deitcher, Edwin S. Levitan

## Abstract

Neuropeptides control rhythmic behaviors, but the timing and location of their release within circuits is unknown. Here imaging in the brain shows that synaptic neuropeptide release by *Drosophila* clock neurons is diurnal, peaking at times of day that were not anticipated by prior electrical and Ca^2+^ data. Furthermore, hours before peak synaptic neuropeptide release, neuropeptide release occurs at the soma, a neuronal compartment that has not been implicated in peptidergic transmission. The timing disparity between release at the soma and terminals results from independent and compartmentalized mechanisms for daily rhythmic release: consistent with conventional electrical activity-triggered synaptic transmission, terminals require Ca^2+^ influx, while somatic neuropeptide release is triggered by the biochemical signal IP_3_. Upon disrupting the somatic mechanism, the rhythm of terminal release and locomotor activity period are unaffected, but the number of flies with rhythmic behavior and sleep-wake balance are reduced. These results support the conclusion that somatic neuropeptide release controls specific features of clock neuron dependent behaviors. Thus, compartment specific mechanisms within individual clock neurons produce temporally and spatially partitioned neuropeptide release to expand the peptidergic connectome underlying daily rhythmic behaviors.

**Significance Statement:** It is believed that electrical activity simultaneously stimulates widespread release sites in single neurons to elicit neuropeptide dependent behaviors. However, optically detecting neuropeptide release in the intact brain shows that clock neurons release neuropeptides from different sites at different times of the day. This is possible because one neuronal compartment, the soma, uses biochemical signaling instead of electrical activity to evoke release. Disrupting somatic release affects specific features of circadian locomotor activity and sleep. Thus, neuropeptide release is elicited by independent triggers from distinct parts of clock neurons to engage different regions of behavior regulating circuitry. This strategy for expanding the connectome may be used for other neuropeptide dependent behaviors, such as feeding and pain perception.

## Introduction

Neuropeptides control development, synaptic function and behavior. Studies of neuropeptide release within neural networks began with immunodetection of neuropeptide content decreases (Taghert and Nitabach, 2012; Nusbaum *et al*., 2017). However, this approach is only applicable when neuropeptide release is dramatic and relies on assuming that transport, capture and degradation of neuropeptide-containing dense-core vesicles (DCVs) do not contribute to measured changes. Given that detecting neuropeptide release at living synapses is challenging, recent studies have measured somatic electrical activity and Ca^2+^ as surrogates for action potential-evoked Ca^2+^ influx that triggers synaptic release. Yet, these indirect approaches do not provide a clear consensus about the timing of neuropeptide release by *Drosophila* small and large ventral lateral (s-LNv and l-LNv) clock neurons, which participate in circadian rhythms and the control of sleep. For example, membrane excitability of s-LNv and l-LNv neurons peaks slowly near sunrise, while cytosolic Ca^2+^ in l-LNv neurons slowly peaks near midday ∼7 hours after the s-LNv neurons (Park *et al*., 2000; Cao & Nitabach, 2008; Sheeba *et al*., 2008; Liang *et al*., 2016). To resolve this discrepancy, we explored whether a recently developed optical approach that detects when and where neuropeptide release occurs at peripheral synapses (Bulgari *et al*., 2019) can be applied to central clock neurons.

Here neuropeptide release by s-LNv and l-LNv clock neurons is resolved in the intact *Drosophila* brain. Synaptic release is found to be rhythmic, but occurs with timing that was not evident from electrical or Ca^2+^ recordings. Even more remarkable, the soma, an unconsidered compartment for release in connectome and physiology studies, is a site of endogenous release that occurs with a different daily schedule than terminals. Genetically disabling the rhythmic somatic release trigger alters features of daily locomotor activity and sleep. Thus, distinct neuronal compartments are demonstrated to use different release mechanisms that in turn control different aspects of rhythmic behavior.

## Results and Discussion

### Timing and Location of Neuropeptide Release by Clock Neurons

We began by considering whether the circadian function of s-LNv and l-LNv neurons is preserved after expressing fluorescent neuropeptide sensors. For *Drosophila* s-LNv neurons, an early attempt at using a GFP-tagged mammalian neuropeptide was not fruitful because of genetic background effects (Kula *et al*., 2006). However, we found that rhythmic changes in the native neuropeptide pigment dispersing factor (PDF) in s-LNv neuron terminals and somas (Park *et al*., 2000) are recapitulated by GFP-tagged *Drosophila* preproinsulin-like peptide 2 (Dilp2-GFP) (Fig. S1A, B). Furthermore, the known absence of rhythmic changes in PDF content of l-LNv neurons was also evident with Dilp2-GFP (Fig. S1C). The preservation of normal function led us to use a more sensitive release indicator, a Dilp2-based construct that employs a fluorogen activating protein (FAP) mL5** with high affinity (Kd ∼20 pM) and a membrane-excluded far-red fluorogenic dye (MG-bTau or tCarb) (Szent-Gyorgyi *et al*., 2013; Yan *et al*., 2015; Pratt *et al*., 2017), which produces fluorescence upon the opening of DCV fusion pores that mediate synaptic release of neuropeptides (Bulgari *et al*., 2019). In l-LNv terminals ongoing release was evident and, as expected, was enhanced with K^+^-induced depolarization and inhibited by chelating extracellular Ca^2+^ with EGTA (Fig. S2). Furthermore, circadian locomotor activity was preserved in animals expressing Dilp2-FAP in LNv neurons (Supplemental Table 1). With the feasibility of FAP-based endogenous release measurements established in the adult brain, this approach was used to measure neuropeptide release across the day in these rhythmically active neurons.

These experiments produced a number of unexpected results. First, data from s-LNv and l-LNv terminals showed that the time of day for peak endogenous synaptic release do not correlate with the timing of electrical activity or Ca^2+^ (see Introduction): LNv neuron synaptic neuropeptide release peaks rapidly 3 hours after sunrise (ZT3) (Fig. 1Ai, ii) and is *preceded* by synaptic release by l-LNv neuron terminals, which occurs in a burst late at night (ZT23) (Fig. 1Bi, ii). These data imply that past activity and Ca^2+^ measurements were not sufficient for inferring the timing of neuropeptide release by clock neuron terminals.

**Figure 1.**
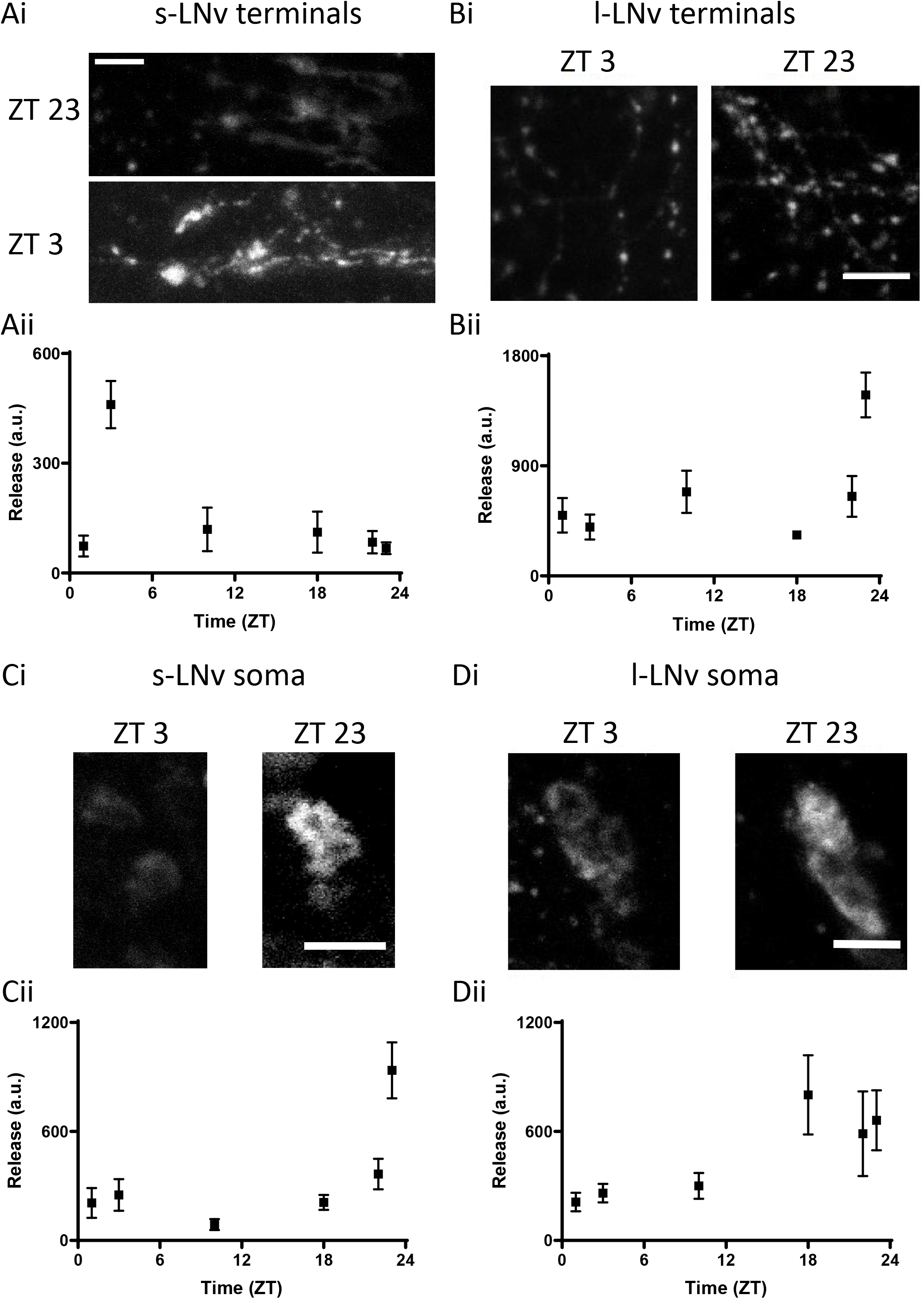
Multi-compartmental neuropeptide release by two subsets of ventrolateral clock neurons exhibits daily rhythms. Ai. FAP images of neuropeptide release at s-LNv terminals at ZT3 than ZT23 in entrained flies (12L:12D). Scale bar = 10 μm. ii. Quantification of neuropeptide release from s-LNv terminals. n = 5-22. Bi. Images of neuropeptide release at l-LNv nerve terminals at ZT3 and ZT23 (scale bar = 10 μm). ii. Quantification of neuropeptide release by l-LNv nerve terminals across the day and night. n = 8-13 for each point. Ci. Imaging shows greater release at s-LNv somas at ZT23 than ZT3 in entrained flies (12L:12D). Scale bar = 10 μm. ii. Quantification of neuropeptide release reveals s-LNv somas secrete neuropeptide across the day and night, with a large spike in release occurring at the end of the night (12L:12D). n=10-18. Di. Images of neuropeptide release in l-LNv somas during the morning (ZT3) and late night (ZT23). Scale bar =10 μm. ii. Neuropeptide release by l-LNv somas across the day and night. n = 6-13 for each time point.

Second, although neuronal somas have not been implicated previously in peptidergic transmission *in vivo*, FAP imaging revealed endogenous somatic neuropeptide release by LNv clock neurons (Fig. 1C, D). Even more remarkable, somatic release occurred hours earlier than release by distal terminals of the same neurons; peak somatic release by s-LNv neurons occurred at ZT23, while l-LNv neurons maintained an elevated release between ZT18 and ZT23 (Fig. 1Cii, Dii). Even though electrical signaling in this subset of neurons is required for normal clock circuit function and behavior (Nitabach *et al*., 2002), the different timing of release from terminals and the soma excludes propagating action potentials as the sole mechanism for evoking neuropeptide release.

### Compartment-Specific Release Mechanisms

Because the effect of action potentials could be gated by other mechanisms, it was possible that electrical activity, which fluctuates slowly during the day, is necessary (but not sufficient) for release. This hypothesis predicts that inhibiting plasma membrane Ca^2+^ channels activated by action potentials should inhibit release throughout the neuron. Therefore, we examined the effect of the Ca^2+^ channel blocker Cd^2+^ on release by each compartment. Peak synaptic neuropeptide release was inhibited, as expected, but, contrary to the hypothesis, somatic neuropeptide release was not affected in either s-LNv or l-LNv neurons (Fig. 2A, B). These data imply that neuropeptide release by the soma, in contrast to release by terminals, does not require Ca^2+^ influx, thus excluding participation of action potentials in somatic release.

**Figure 2.**
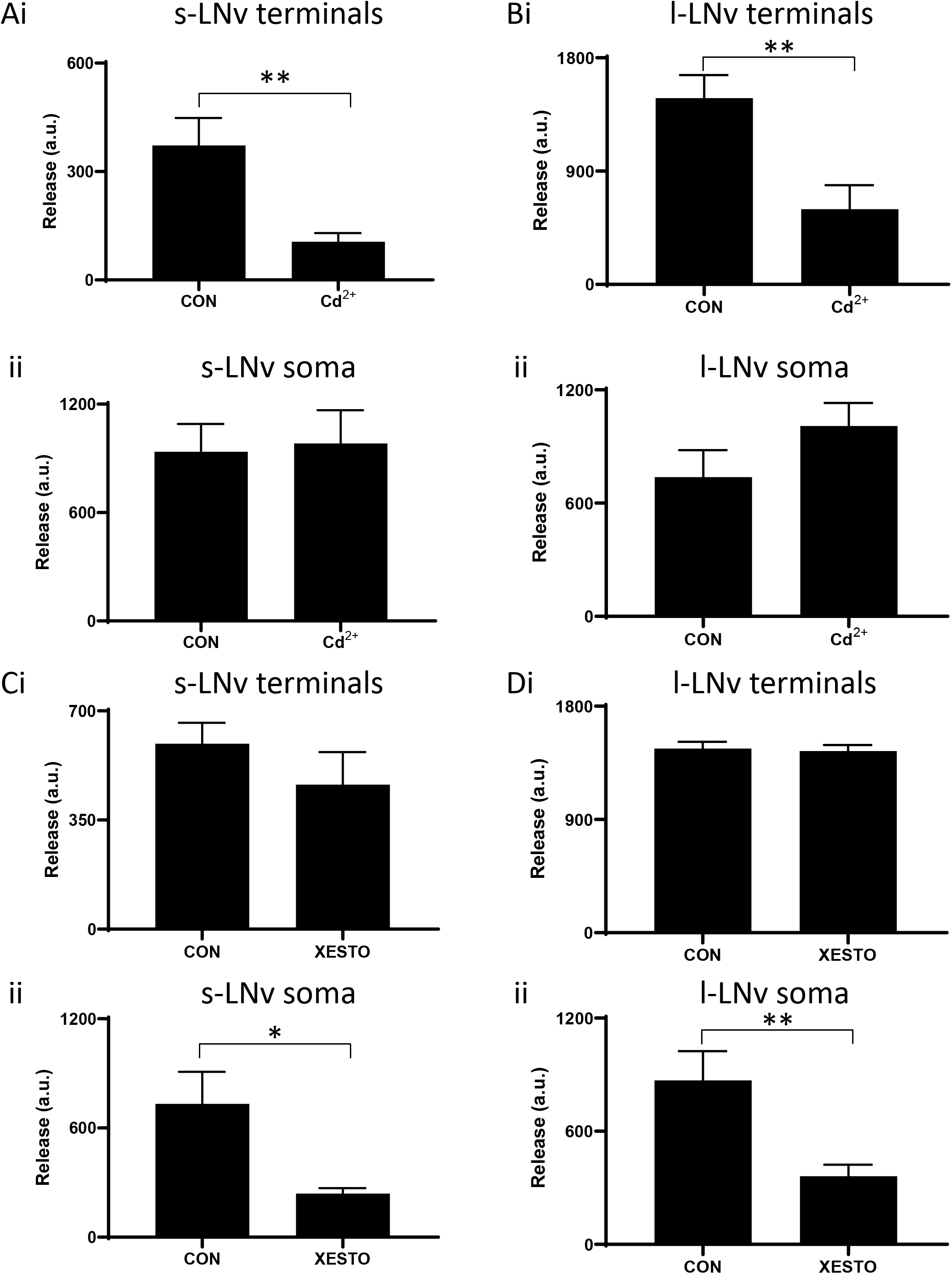
Peak neuropeptide release requires Ca^2+^ influx into terminals and IP_3_ signaling in the soma. Ai. Peak release (ZT3) by s-LNv terminals is inhibited by Cd^2+^ (10 μM, n = 13) compared to controls (n = 14). **p < 0.01, unpaired t test. ii. Peak release (ZT 23) by s-LNv somas is not affected by Cd^2+^ (10 μM, n = 13). For CON, n = 14). P = 0.8493, unpaired t test. Bi. Peak release (ZT23) by l-LNv terminals is inhibited by Cd^2+^ (10 μM, n = 10) compared to controls (n = 8). **p < 0.01, unpaired t test. ii. Peak release (ZT 23) by l-LNv somas is not affected by Cd^2+^ (10 μM, n = 10). For CON, n = 8). P = 0.8493, unpaired t test. Ci. Peak release (ZT3) by s-LNv terminals is not different between Xestospongin C (20 μM, n = 9) and control terminals (CON, n = 12). p = 0.7951, unpaired t test. ii. Peak release (ZT23) by s-LNv somas is significantly reduced by Xestospongin C (20 μM) (n = 19) compared to controls (CON, n = 24). *p < 0.05, unpaired t test. Di. Peak release (ZT23) by l-LNv terminals is not different between Xestospongin C (20μM, n = 9) and controls (CON, n = 16). p = 0.3126, unpaired t test. ii. Peak release (ZT 23) by l-LNv somas is reduced by Xestospongin C (20 μM) (n = 16) compared to controls (CON, n = 21). **p < 0.01, unpaired t test.

If electrical activity does not couple release from terminals and the soma, what mechanism is responsible for somatic neuropeptide release? Recently, FAP imaging demonstrated that there is spontaneous neuropeptide release, which is independent of extracellular Ca^2+^ and, in contrast to activity-dependent release, resistant to tetanus toxin (TeTx) (Bulgari *et al*., 2019). However, TeTx expression was found to inhibit neuropeptide release by s-LNv and l-LNv somas (Fig. S3). The incomplete inhibition of release may explain why TeTx fails to disrupt circadian behaviors mediated by the neuropeptide PDF, which is an important clock signal released by s-LNv and l-LNv neurons (Kaneko *et al*., 2005; Blanchardon *et al*., 2001). More relevant here, TeTx inhibition suggests that somatic neuropeptide release is Ca^2+^ dependent.

The implication of Ca^2+^ dependence (Fig. S3) without extracellular Ca^2+^ influx (Fig. 2A, B) led us to examine the role of IP_3_ (inositol trisphosphate) receptors (IP3Rs), which mediate intracellular Ca^2+^ release from the endoplasmic reticulum into the cytosol. First, 100 nM Xestospongin C, a blocker of IP3Rs (Gafni *et al*., 1997; Klose *et al*., 2010; James *et al*., 2019), was applied acutely. Strikingly, while peak endogenous neuropeptide release from terminals was unaffected, peak endogenous neuropeptide release from the soma was inhibited (Fig. 2C, D). To independently test for a role for IP_3_ signaling in somatic neuropeptide release, IP_3_ sponge, which binds IP_3_ to interfere with its second messenger function (Uchiyama *et al*., 2002; Usui-Aoki *et al*., 2005; James *et al*., 2019), was expressed along with the FAP reporter in LNv clock neurons and tested for effects on neuropeptide release at times of the day when release varies in terminals and the soma (ZT23 and ZT3, Fig. 1). Consistent with the Xestospongin C results, neuropeptide release from LNv terminals expressing IP_3_ sponge remained rhythmic (Fig. 3Ai, Bi) and peak somatic release at ZT23 was abolished (Fig. 3Aii, Bii), thus showing that the somatic release rhythm was disrupted. These data show that neuropeptide release by terminals does not depend on preceding somatic neuropeptide release. Furthermore, different neuronal compartments release neuropeptides at different times of the day by engaging different triggers (i.e., extracellular Ca^2+^ influx for terminals and IP_3_ signaling for the soma).

**Figure 3.**
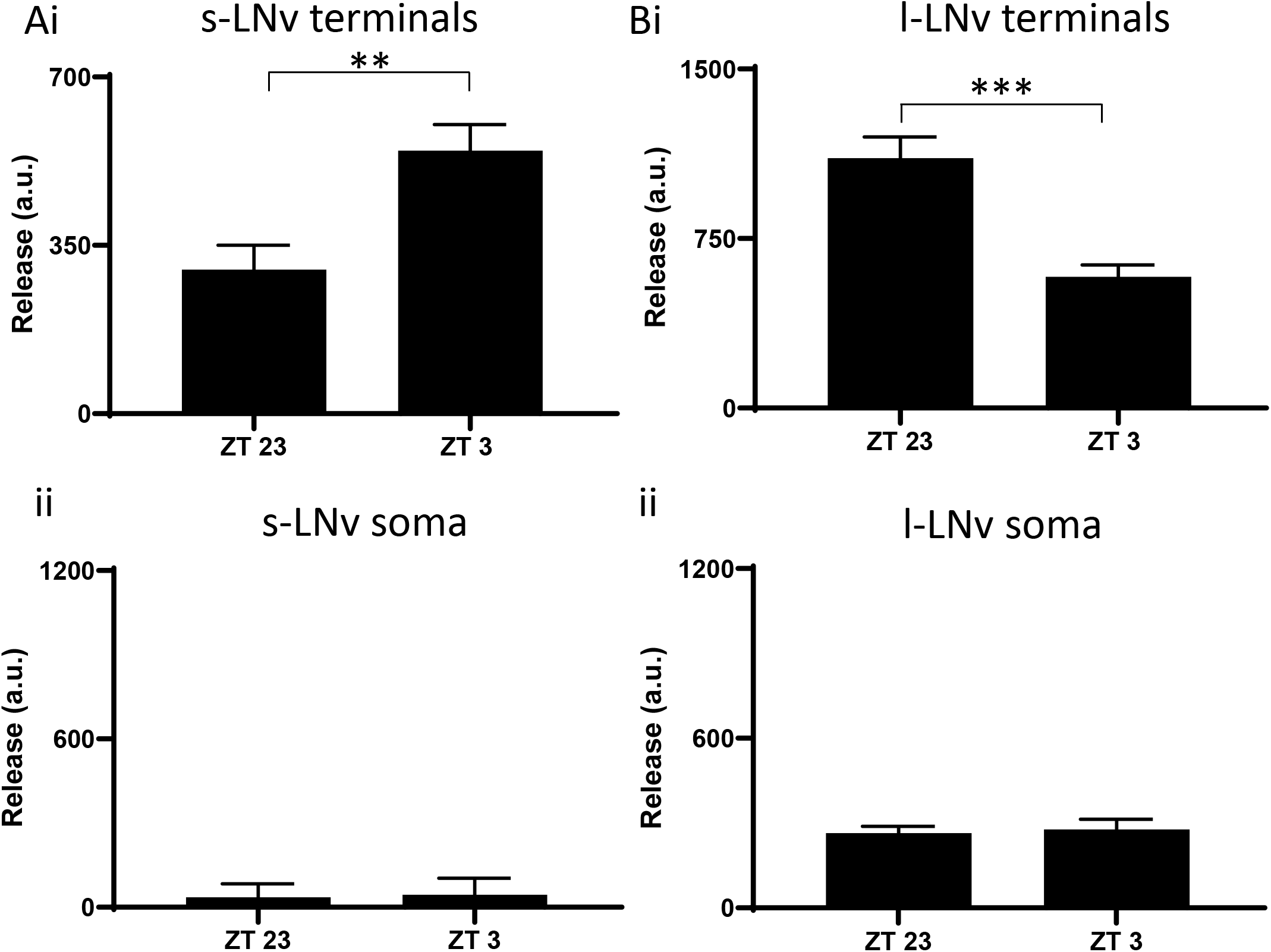
IP_3_ sponge disrupts rhythmic somatic neuropeptide release in LNv clock neurons. Ai. Circadian rhythms in s-LNv terminal neuropeptide release continued when IP_3_ signaling is disrupted through expression of IP_3_ sponge in PDF cells. Note the difference between ZT23 (n = 13) and ZT3 (n = 13), **p < 0.01, unpaired t test. i. Neuropeptide release from s-LNv somas is abolished when IP_3_ signaling is disrupted through expression of IP_3_ sponge in PDF cells. Note that there was no significant difference between ZT23 (n=8) and ZT3 (n = 7), p = 0.907, unpaired t test. Bi. Circadian rhythms in l-LNv terminal neuropeptide release continued when IP_3_signaling is disrupted through expression of IP_3_ sponge in PDF cells. Note the difference between ZT23 (n = 6) and ZT3 (n = 6), ***p < 0.001, unpaired t test. ii. Neuropeptide release from l-LNv somas is interrupted when IP_3_ signaling is disrupted through expression of *IP*_*3*_ *sponge* in PDF cells. Note that there was no significant difference between ZT23 (n = 6) and ZT3 (n = 6), p = 0.7690, unpaired t test.

### Behavioral Effects of Compartmental Release Mechanisms

Transmission by the neuropeptide PDF from s-LNv neurons regulates several aspects of circadian behavior including morning anticipation, free-running rhythm period and rhythmicity (Renn *et al*., 1999; Hyun *et al*. 2005; Choi *et al*., 2012; Klose *et al*., 2016). Because IP_3_ sponge selectively ablates somatic neuropeptide release, we examined the behavior of flies expressing IP_3_ sponge in LNv neurons for effects on clock function behavioral output (Fig. 4A-C). Expression of IP_3_ sponge with Dilp2-FAP in LNv neurons did not affect morning anticipation or the period of daily locomotor rhythms, but reduced the number of flies that maintain rhythmic locomotor behavior compared to genetic controls, and increased their daily activity (Fig. 4D).

**Figure 4.**
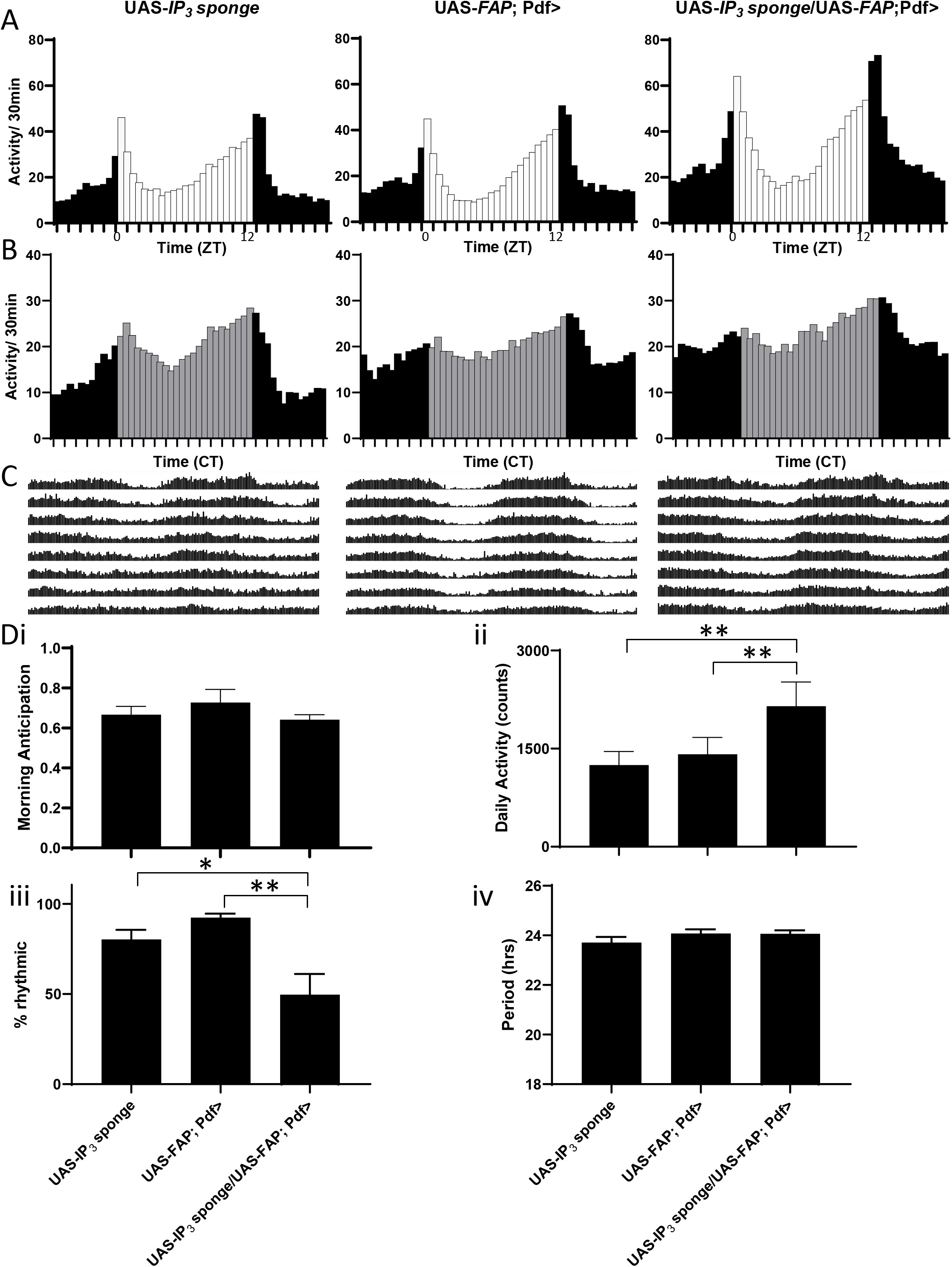
IP_3_ signaling in LNv neurons suppresses locomotor activity and promotes rhythmicity. A. LD group eductions for progeny of UAS-*FAP*; Pdf-Gal4 flies crossed to UAS-IP_3_ sponge (n = 32), UAS-FAP; Pdf-Gal4 (n = 32), UAS-IP_3_ sponge (n = 32), are shown. 12h light (white bars):12h dark (black bars). B. DD group eductions. Constant darkness (grey bars indicate subjective day). C. Average group actograms displayed in double plotted format over DD2-8. D. Behavioral indices for each genotype including: i. morning anticipation (repeated measures (RM) one-way ANOVA revealed no significant difference (p = 0.1743), ii. total locomotor activity (RM one-way ANOVA revealed significant difference (p < 0.01)), iii. Rhythmicity (RM one-way ANOVA revealed significant difference (p < 0.01)), and period (RM one-way ANOVA revealed no significant difference (p 0.3894). *p < 0.05, **p < 0.01, Dunnett’s multi-comparison test

The latter change (Fig. 4Dii) led us to consider whether sleep was affected. Quantification of sleep behavior revealed that the increase in overall activity was associated with reduced total sleep time and a large increase in the wake/sleep ratio (Fig. 5A-D). Interestingly, average length of individual sleep bouts was shortened without a change in sleep bout number, thus revealing an effect on sleep consolidation (Fig. 5E, F). Together with the circadian locomotor activity studies, these experiments identified numerous behavioral parameters (circadian period, morning anticipation and sleep bout frequency) that remain normal when somatic release is inhibited. Thus, for those behaviors synaptic neuropeptide release by terminals is sufficient. However, the experimental results also show that a different subset of behavioral parameters (circadian rhythmicity and sleep consolidation) rely on an action potential independent mechanism that supports somatic neuropeptide release.

**Figure 5.**
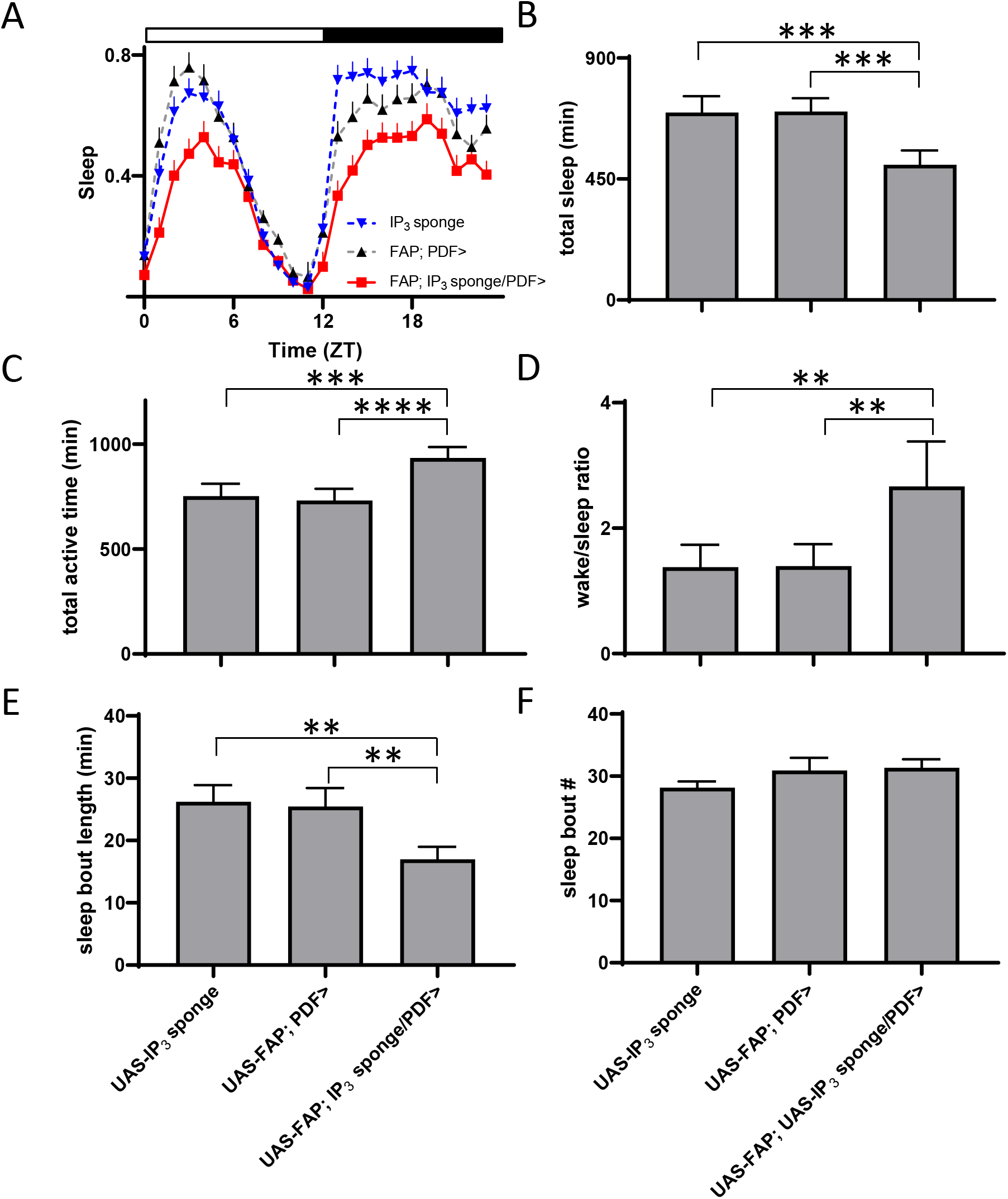
IP_3_ signaling in LNv neurons increases total sleep time through promotion of sleep consolidation. A. Sleep decreases in flies expressing IP_3_ sponge in PDF neurons (UAS*-*IP_3_ sponge/UAS*-*FAP; Pdf-Gal4) compared to two genetic controls (UAS*-*IP_3_ sponge and UAS*-FAP*; *Pdf-Gal4)*. Sleep presented as fraction of each hour across the day. Flies entrained under a 12h:12-h light/dark (LD) regimen. Data shown are from one experiment with 25 flies in each genotype. B. Sponge expression in PDF neurons decreases total sleep by over three hours compared to controls. A Repeated Measures (RM) one-way ANOVA revealed significant difference (p < 0.001). C. Total active time. A RM one-way ANOVA revealed significant difference (p < 0.001). D. Sleep/wake ratio. A RM one-way ANOVA revealed significant difference (p < 0.001). E. Sleep bout length. A RM one-way ANOVA revealed significant difference (p < 0.01). F. Sleep bout number. A RM one-way ANOVA revealed no significant difference (p = 0.1816). Data analyzed from averages of five experiments. Total fly counts for each genotype: UAS-IP_3_ sponge/UAS*-*FAP; Pdf-Gal4 (n = 161), UAS*-*IP_3_ sponge (n = 164) and UAS*-* FAP; Pdf-Gal4 (n = 167). *** indicates p < 0.001, ** indicates p < 0.01, Dunnett’s multiple-comparison test.

FAP imaging revealed synaptic neuropeptide release from LNv clock neurons that does not conform to predictions from previously used indirect methods. Content measurements could not resolve whether somatic changes were due to release or traffic, and also did not detect l-LNv rhythmic neuropeptide release, likely because it is relatively modest and/or obscured by DCV capture that replenishes synaptic neuropeptide stores (Shakiryanova *et al*., 2006; Wong *et al*., 2012). Furthermore, somatic Ca^2+^ was not reflective of release at terminals likely because of somatic IP_3_ signaling. Finally, somatic electrical recording cannot take into account regulation by presynaptic inputs. Thus, direct live imaging of neuropeptide release is essential for monitoring peptidergic transmission in the brain.

Indeed, this approach demonstrates that that central clock neurons release neuropeptide from terminals and the soma, with each compartment operating with different mechanisms and timing. Release from LNv clock neuron terminals is conventional (i.e., mediated by extracellular Ca^2+^ influx), but somatic neuropeptide release is triggered by IP_3_ signaling that operates in the absence of action potential-induced Ca^2+^ influx. Different release mechanisms allow for multi-phasic temporal control of neuropeptide release from separate compartments of the same neuron, each of which releases onto different parts of the clock circuit, thereby providing separate output avenues to independently influence different parameters of behavior.

## Materials and Methods

### Flies

All flies were reared on cornmeal/agar supplemented with yeast. Male flies were collected on the day of eclosion and maintained on a 12 hour light: 12 hour dark photoperiod for physiological experiments. Flies were maintained at 25°C. Genotypes include: UAS-*Dilp2-GFP*; *Pdf*-Gal4, UAS-*Dilp2-FAP*; *Pdf*-Gal4, UAS-*Dilp2-FAP*/UAS-*Dilp2-GFP*; *Pdf*-Gal4, UAS-*Dilp2-FAP*; UAS-*IP*_*3*_ *sponge Pdf*-Gal4, UAS-*Dilp2-FAP*; UAS-*IP*_*3*_ *sponge*. UAS-*Dilp2-GFP*, UAS-*Dilp2-FAP* and UAS-*IP*_*3*_ *sponge* were described previously (Wong *et al*., 2012; Usui-Aoki *et al*., 2005; Bulgari *et al*., 2019). Expression of tetanus toxin light chain and a control mutant (Sweeney *et al*., 1995) were induced by crosses to UAS lines (Bloomington #28838 and 28840).

### Imaging

Experiments were performed on adult brain explants, bathed in HL3 saline, which contained (in mM) 70 mM NaCl, 5 KCl, 1.5 CaCl_2_, 20 MgCl_2_, 10 NaHCO_3_, 5 trehalose, 115 sucrose, and 5 sodium Hepes, pH 7.2. Brains were dissected in Ca^2+^-free HL3 saline (0.5 mM EGTA in place of Ca^2+^), transferred to HL3 saline and then placed in HL3 filled Sylgard chambers that were pre-treated with poly-L-lysine, which acted as an adherent. To elicit depolarization-evoked release, HL3 was modified by increasing KCl to 70 mM by substituting NaCl.

Imaging data were acquired with an Olympus Fluoview 1000 upright confocal microscope with a 60× 1.0 NA water immersion objective. FAP signals were imaged with 640 nm excitation and Cy5 fluorescence optics, while GFP was imaged with a 473 nm excitation laser and standard FITC optics. For FAP experiments, membrane impermeant fluorogen (MG-B-Tau (Figure 1) or MG-TCarb (Figure 2-5)) was added to the bathing solution at a final concentration of 1 µM. Endogenous release measurements (Figure 2-5) were acquired after 60 minutes of bathing in the fluorogens.

Quantification of fluorescence intensity was performed with ImageJ software (https://imagej.nih.gov/ij/). For s-LNv terminal GFP content, a region of interest was drawn around the s-LNv dorsal protocerebral projection stack. l-LNv GFP terminal content was the average of 10 boutons for each hemi-segment. For s-LNv terminal release, 6-10 boutons were averaged for each measurement. s-LNv and l-LNv soma measurements are the average of 2-4 cells from each hemi-segment. Images were taken at several times points during both the day and night and all measurements were background subtracted.

### Behavior

Adult flies (3-4 days old) were loaded into glass tubes and placed in DAM2 Trikinetics Activity Monitors and entrained for 7 days on 12hr:12hr light:dark schedule, then released into constant darkness for 8 days. We assessed rhythmicity by normalizing activity from DD days 2–8. We defined arrhythmic flies by rhythmicity threshold [Qp.act/Qp.sig] below 1 or a period estimate <18 hr or >30 hr. Rhythmicity and period were assessed using ShinyR software (Cichewicz and Hirsh, 2018).

### Analysis

Statistical analysis and graphing were performed with Graphpad Prism software. Error bars represent Standard Error of the Mean (SEM). Statistical comparison for two experimental groups was based on Student’s t test. For multiple comparisons, one-way ANOVA was followed with Dunnett’s post-test.

### Chemicals

MG-B-tau and MG-TCarb were synthesized as previously described (Yan *et al*., 2015; Pratt *et al*., 2017). Pharmacological agents were bath applied in recording saline. Purchased chemicals included Xestospongin C (CAS Number: Abcam 88903-69-9) and cadmium chloride (SigmaAldrich).

## Acknowledgements

We thank Dmytro Kolodieznyi (Carnegie Mellon University) for the preparation and characterization of the MG-TCarb dye, and C. Andrew Frank (University of Iowa) for providing UAS-*IP*_*3*_ *sponge* flies. This research was supported by NIH grants R01NS32385 and R21NS11523 to ESL and RF1MH115023 and R21MH100612 to MPB.

## Legends

**Figure S1.**
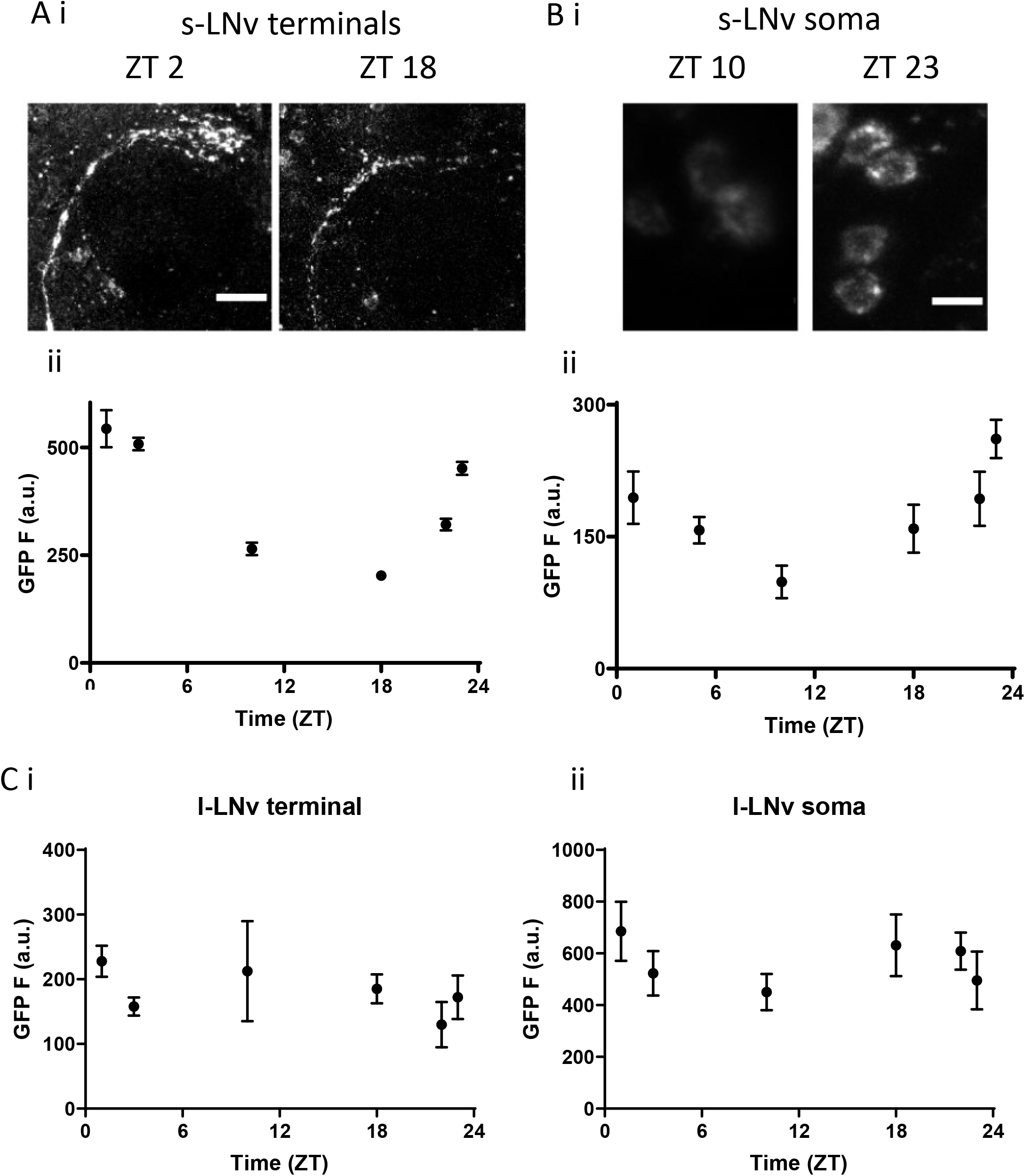
Neuropeptide imaging of LNv neurons in intact living brains with Dilp2-GFP. A. i. Dilp2-GFP images of s-LNv terminal projections into the dorsal protocerebrum (Z-projection stack) at ZT 2 and ZT 18. Scale bar = 25 μm. ii. Quantification of neuropeptide content from s-LNv terminals. n = 5-22 B. i. Dilp2-GFP images of s-LNv somas at ZT2 and ZT23. Scale bar = 10 μm. ii. Quantification of neuropeptide content from s-LNv somata. n = 10-18 C. i. l-LNv terminal neuropeptide content assessed at 6 time points in a 24-hour period in flies that were entrained 12h light:12h dark. n = 7-13 for each time point. ii. l-LNv s omatic neuropeptide content assessed at 6 time points over 24 hours in flies that were entrained 12h light:12h dark. n = 6-12 for each time point.

**Fig. S2.**
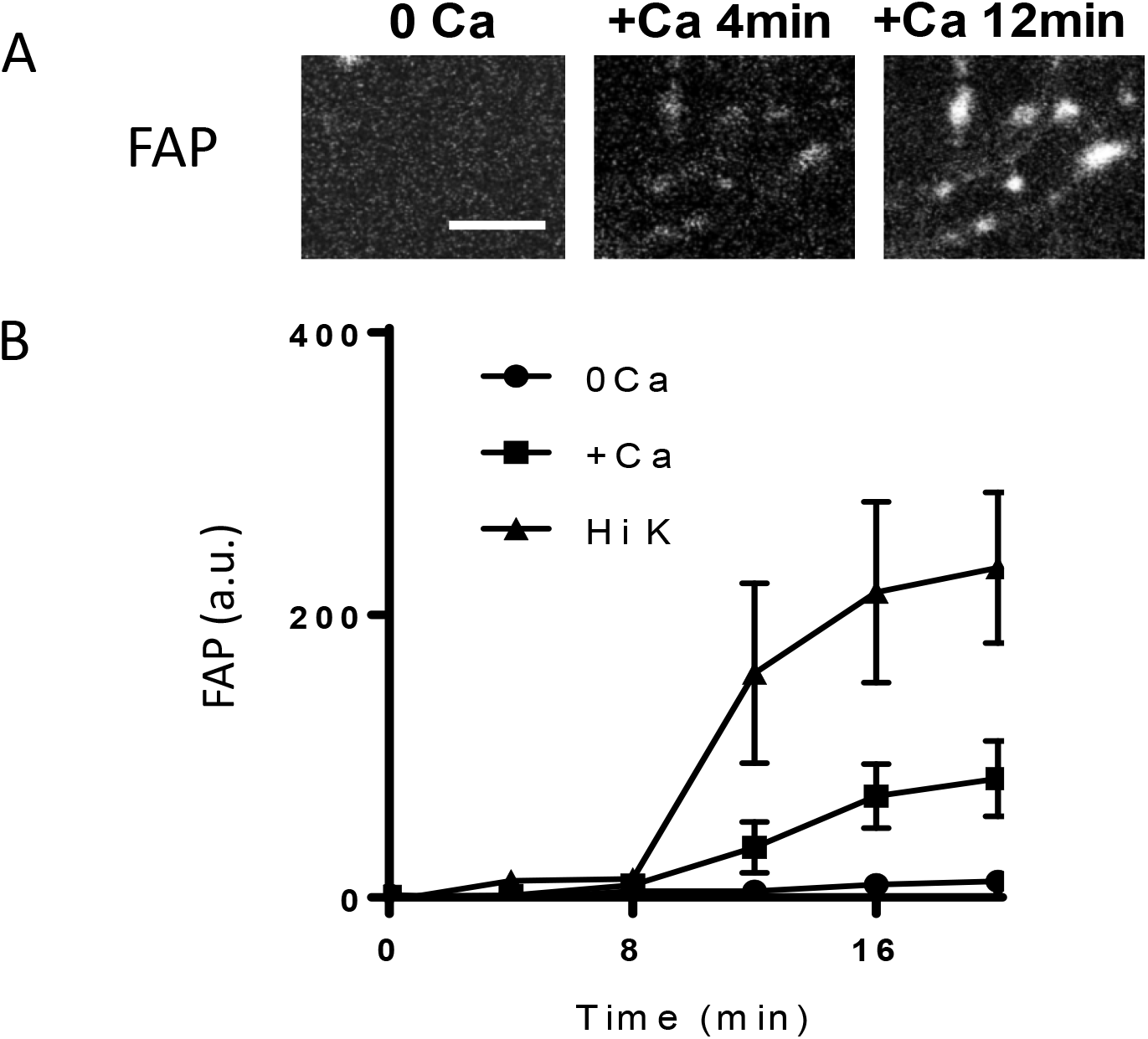
Neuropeptide release from l-LNv terminals. A. Dilp2-FAP (FAP) in l-LNv nerve terminals reveals peptide release. Excised brains started in 0 Ca^2+^. After 8 min the bath was exchanged with Ca^2+^ containing saline (+Ca) to measure the effect of intrinsic activity. Note that +Ca increases the FAP signal within 4 min. Bar, 5M µm. B. FAP quantification comparing 0 Ca, +Ca and depolarization with high K^+^ in the presence of Ca^2+^ (Hi K). n > 3.

**Fig. S3.**
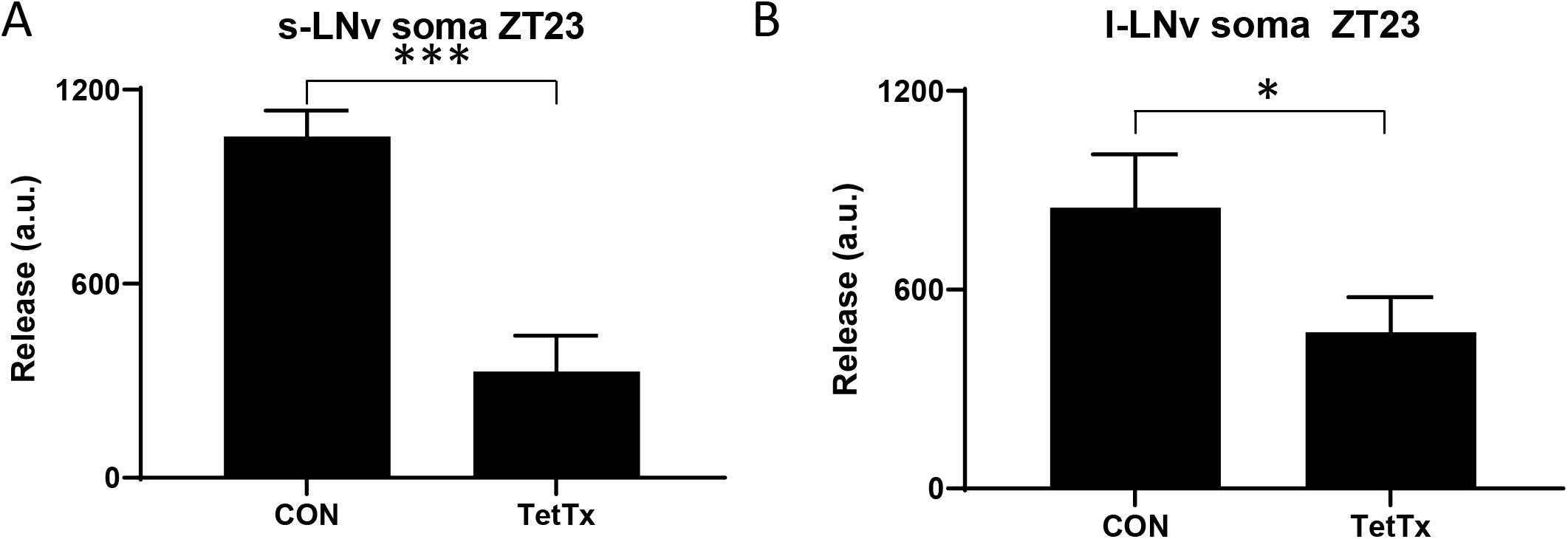
Tetanus Toxin (TetTx) attenuates neuropeptide release from LNv somas. A. Release from s-LNv somas at ZT23 expressing tetanus toxin (n = 12) was 69% less than controls (n = 8) expressing inactive mutant toxin. ***p < 0.001, unpaired t test. B. Release from l-LNv somas at ZT23 expressing tetanus toxin (n = 22) was 44% less than controls (n = 14) expressing inactive mutant toxin. *p < 0.05, unpaired t test.

**Supplemental Table. 1.** Rhythmic Locomotor Activity in days DD2-DD8 for flies expressing IP_3_ sponge to alter neuropeptide release. Chi-squared period testing range was 18-30h, resolution was 0.2h, and threshold for filtering arrhythmic flies was set at a rhythm strength of 1, thus any flies below this threshold were not considered rhythmic and did not contribute to the calculation of period. n = # of flies; N = # of experiments; AR% = % of arrhythmic flies; Tau = period of rhythmic flies.

|  | n | N | TAU | SEM | %AR | Rhythm strength | SEM |
| --- | --- | --- | --- | --- | --- | --- | --- |
| UAS-IP3 sponge | 120 | 5 | 23.71 | 0.23 | 7 | 1.35 | 0.04 |
| UAS-FAP; Pdf> | 107 | 5 | 24.08 | 0.16 | 19 | 1.34 | 0.03 |
| UAS-FAP; Pdf><br>/UAS-IP3 sponge | 120 | 5 | 24.06 | 0.14 | 50 | 1.11 | 0.01 |

## Notes

### Competing Interest Statement

The authors have declared no competing interest.

### Summary of Updates

New data including changes in sleep

## References

Blanchardon E, Grima B, Klarsfeld A, et al. Defining the role of Drosophila lateral neurons in the control of circadian rhythms in motor activity and eclosion by targeted genetic ablation and PERIOD protein overexpression. Eur J Neurosci. 2001;13(5):871–888.

Bulgari D, Deitcher DL, Schmidt BF, Carpenter MA, Szent-Gyorgyi C, Bruchez MP, Levitan ES. Activity-evoked and spontaneous opening of synaptic fusion pores. Proc Natl Acad Sci U S A. (2019) 116(34):17039–17044.

Cao G, Nitabach MN. Circadian control of membrane excitability in Drosophila melanogaster lateral ventral clock neurons. J Neurosci. 2008;28(25):6493–6501.

Choi C, Cao G, Tanenhaus AK, et al. Autoreceptor control of peptide/neurotransmitter corelease from PDF neurons determines allocation of circadian activity in drosophila. Cell Rep. 2012;2(2):332–344.

Cichewicz K. & Hirsh J., ShinyR-DAM: A program analyzing Drosophila activity, sleep, and circadian rhythms, Commun. Biol. 2018; 1, 25.

Gafni J, Munsch JA, Lam TH, Catlin MC, Costa LG, Molinski TF, Pessah IN. Xestospongins: potent membrane permeable blockers of the inositol 1,4,5-trisphosphate receptor. Neuron. 1997; 19:723–733.

Hyun S, Lee Y, Hong ST, et al. Drosophila GPCR Han is a receptor for the circadian clock neuropeptide PDF. Neuron. 2005;48(2):267–278.

James TD, Zwiefelhofer DJ, Frank CA. Maintenance of homeostatic plasticity at the Drosophila neuromuscular synapse requires continuous IP3-directed signaling. Elife. 2019;8:e39643.

Kaneko M, Park JH, Cheng Y, Hardin PE, Hall JC. Disruption of synaptic transmission or clock-gene-product oscillations in circadian pacemaker cells of Drosophila cause abnormal behavioral rhythms. J Neurobiol. 2000;43(3):207–233.

Klose MK, Dason JS, Atwood HL, Boulianne GL, Mercier AJ. Peptide-induced modulation of synaptic transmission and escape response in Drosophila requires two G-protein-coupled receptors. J Neurosci. 2010;30(44):14724–14734.

Klose M, Duvall L, Li W, et al. Functional PDF Signaling in the Drosophila Circadian Neural Circuit Is Gated by Ral A-Dependent Modulation. Neuron. 2016;90(4):781–794.

Kula E, Levitan ES, Pyza E, Rosbash M. PDF cycling in the dorsal protocerebrum of the Drosophila brain is not necessary for circadian clock function. J Biol Rhythms. 2006; 21(2):104–117.

Liang X, Holy TE, Taghert PH. Synchronous Drosophila circadian pacemakers display nonsynchronous Ca^2^+ rhythms in vivo. Science. 2016;351(6276):976–981.

Nitabach MN, Blau J, Holmes TC. Electrical silencing of Drosophila pacemaker neurons stops the free-running circadian clock. Cell. 2002;109(4):485–495.

Nusbaum MP, Blitz DM, Marder E, Functional consequences of neuropeptide and small-molecule co-transmission. Nat. Rev. Neurosci. 18, 389–403 (2017).

Park JH, Helfrich-Förster C, Lee G, Liu L, Rosbash M, Hall JC. Differential regulation of circadian pacemaker output by separate clock genes in Drosophila. Proc Natl Acad Sci U S A. 2000;97(7):3608–3613.

Pratt CP, Kuljis DA, Homanics GE, He J, Kolodieznyi D, Dudem S, Hollywood MA, Barth AL, Bruchez MP. Tagging of Endogenous BK Channels with a Fluorogen-Activating Peptide Reveals β4-Mediated Control of Channel Clustering in Cerebellum. Front Cell Neurosci. 2017;11:337

Renn SC, Park JH, Rosbash M, Hall JC, Taghert PH. A pdf neuropeptide gene mutation and ablation of PDF neurons each cause severe abnormalities of behavioral circadian rhythms in Drosophila. Cell. 1999;99(7):791–802.

Shakiryanova D, Tully A, Levitan ES. Activity-dependent synaptic capture of transiting peptidergic vesicles. Nat Neurosci. 2006;9(7):896–900.

Sheeba V, Gu H, Sharma VK, O’Dowd D, Holmes TC. Circadian- and Light-Dependent Regulation of Resting Membrane Potential and Spontaneous Action Potential Firing of Drosophila Circadian Pacemaker Neurons. J Neurophysiol. 2008; 99(2): 976–988.

Sweeney ST, Broadie K, Keane J, Niemann H, O’Kane CJ. Targeted expression of tetanus toxin light chain in Drosophila specifically eliminates synaptic transmission and causes behavioral defects. Neuron 14, 341–351 (1995).

Szent-Gyorgyi C, Stanfield RL, Andreko S, Dempsey A, Ahmed M, Capek S, Waggoner AS, Wilson IA, Bruchez MP. Malachite green mediates homodimerization of antibody VL domains to form a fluorescent ternary complex with singular symmetric interfaces. J Mol Biol. 2013;425(22):4595–613.

Taghert, PH, Nitabach MP, Peptide neuromodulation in invertebrate model systems. Neuron 76, 82–97 (2012).

Uchiyama T, Yoshikawa F, Hishida A, Furuichi T, Mikoshiba K. A novel recombinant hyperaffinity inositol 1,4,5-trisphosphate (IP(3)) absorbent traps IP(3), resulting in specific inhibition of IP(3)-mediated calcium signaling. J Biol Chem. 2002;277(10):8106–13.

Usui-Aoki K, et al. Targeted expression of Ip3 sponge and Ip3 dsRNA impaires sugar taste sensation in Drosophila. J Neurogen. 2005; 19:123–141.

Wong MY, Zhou C, Shakiryanova D, Lloyd TE, Deitcher DL, Levitan ES. Neuropeptide delivery to synapses by long-range vesicle circulation and sporadic capture. Cell. 2012;148(5):1029–1038.

Yan Q et al. Near-instant surface-selective fluorogenic protein quantification using sulfonated triarylmethane dyes and fluorogen activating proteins. Org Biomol Chem. 2015; 13(7): 2078–2086.

